# Yeast Dhx29 promotes translation progression by unwinding structured mRNA in the ribosomal A-site

**DOI:** 10.64898/2026.08.24.746666

**Authors:** Leona Chitoiu, Timo Denk, Martin B. D. Müller, Otto Berninghausen, Thomas Becker, Matthias Thoms, Roland Beckmann

## Abstract

mRNAs can form stable structures that need to be resolved to facilitate translation. During translation initiation in mammals, the scanning 48S complex requires the helicase activity of DHX29 to unwind stable mRNA structures that cannot be resolved by eIF4A. Here, we show that the yeast DHX29 homolog, Ylr419w (Dhx29), has a similar function during translation on elongating 80S ribosomes. Cryo-EM analyses show that the Dhx29 helicase module is positioned at the mRNA entry channel to engage mRNA, while its double-stranded RNA-binding domain (dsRBD) senses hairpin-forming mRNA in the ribosomal A-site. By selective ribosome profiling, we observed that Dhx29 is associated with transcripts that form RNA structures, such as stable tetraloops. Dhx29 mutants with perturbed helicase activity enrich 80S with hairpins in the A-site, as well as ribosome collisions, while a mutant lacking the N-terminal dsRBD sensor domain loses the specificity for such ribosomes. We thus propose that Dhx29 functions in translation elongation by resolving structured mRNA formed in the ribosomal A-site through its 3’-5’ helicase activity and pulling on the mRNA from its 3’ end.

## Introduction

Translation is a dynamic process which is safeguarded by various factors that ensure overall efficiency and rapid adaptation to exogenous stimuli. When individual ribosomes stall due to erroneous mRNAs or cellular stress, trailing ribosomes can lead to ribosomal collisions. These collisions trigger stress signaling and ribosome quality control mechanisms, resulting in decay of the problematic mRNA and arrested nascent peptides, as well as the recycling of stalled ribosomes (Inada, 2026; Müller *et al*, 2025). Several ATP-dependent 3’-5’ helicases were shown to engage with 3’-mRNA overhangs, such as Ski2/SKI2 for mRNA degradation (Kögel *et al*, 2024; Müller *et al*., 2025; Schmidt *et al*, 2016; Tomecki *et al*, 2023) or Slh1/ASCC3 (Best *et al*, 2023; Juszkiewicz *et al*, 2020; Matsuo *et al*, 2017; Narita *et al*, 2022) and *E. coli* HrpA for ribosome splitting (Campbell *et al*, 2025). Recently, Ylr419w, another member of the DExH-box helicase family, was shown to be associated with 80S ribosomes and polysomes in the yeast *Saccharomyces cerevisiae* (Fromont-Racine *et al*, 2024). This helicase is closely related to, and was suggested to be, the functional homolog of human DHX29. Initially, DHX29 was described in mammalian cells to promote the formation and scanning of translation initiation complexes on mRNAs with secondary structures in their 5’-UTRs (Hashem *et al*, 2013; Parsyan *et al*, 2009; Pisareva & Pisarev, 2016; Pisareva *et al*, 2008). However, a recent study suggested that it can also engage elongating 80S ribosomes to mediate decay of non-optimal mRNAs (Hia *et al*, 2026). AlphaFold (AF) predictions (Fig. EV1) reveal a similar structural composition of yeast Ylr419w compared to human DHX29 (Dhote *et al*, 2012). This includes a conserved helicase module within its C-terminal part comprising two RecA domains (RecA1 and RecA2), a winged helix (WH), and a ratchet domain, followed by an OB-fold domain. Its N-terminus consists of a double-stranded RNA binding domain (dsRBD) and a UBA-like domain that is partly similar to the DHX29 N-terminal domain (Fig. EV1A-C). In addition, Ylr419w possesses a combined Ubiquitin-Associated (UBA)/RWD domain (named after a conserved domain common in **R**ing-finger proteins, **W**D-repeat proteins, and **D**EAD-like helicases) that is connected to the RecA1 domain via an unstructured linker. Yet, to date, no structural data in context of the translation machinery exist and functional data are limited to mere ribosome binding (Fromont- Racine *et al*., 2024). Here, we report *ex vivo* cryo-EM structures of Ylr419w bound to translating ribosomes, suggesting a function in unwinding mRNA secondary structure blocks forming in the ribosomal A-site. Selective ribosome profiling (ribo-seq) confirmed that Ylr419w functions on 80S ribosomes that harbor mRNA tetraloops, which have a high propensity to form stable RNA structures (Varani, 1995). We thus suggest that Ylr419w plays a role in co-translational unwinding of mRNA hairpin structures forming in the ribosomal A-site that otherwise impede ribosome progression. Our data thus confirms that Ylr419w is a functional homolog of human DHX29, as suggested before (Fromont-Racine *et al*., 2024), and we therefore rename this factor to yeast Dhx29 (Dhx29p).

## Results

### Dhx29 associates with translating ribosomes at the mRNA entry

To investigate ribosome-association of Dhx29, we performed native pulldowns using C-terminally FTpA-tagged (Flag - TEV protease cleavage site - protein A) protein as bait. This approach yielded Dhx29 co-eluting with 80S ribosomes (Fig. 1A). Cryo-EM analysis of this sample revealed that particles predominantly belong to translating 80S ribosomes with mRNA, tRNAs and a nascent polypeptide chain present. However, no 48S initiation complexes were observed (Appendix Fig. S1A). The 80S ribosomes were either in the PRE-translocational state (with tRNAs present in hybrid A/P- and P/E- sites; PRE/hybrid state) or in the POST-translocational state (with tRNA in the P/P site; POST state). The vast majority (about 75%) of these 80S showed extra density emerging from the mRNA entry site of the 40S subunit, representing Dhx29 (Fig. 1C,D and Appendix Fig. S1). In both PRE/hybrid and POST states, Dhx29 is located above the 40S head, spanning from the 40S body (18S rRNA helix h16) along the 40S head towards its beak. For the POST state, additional density for the N-terminus of Dhx29 reaches into the A-site of the 40S subunit (Fig. 1C).

**Fig. 1:**
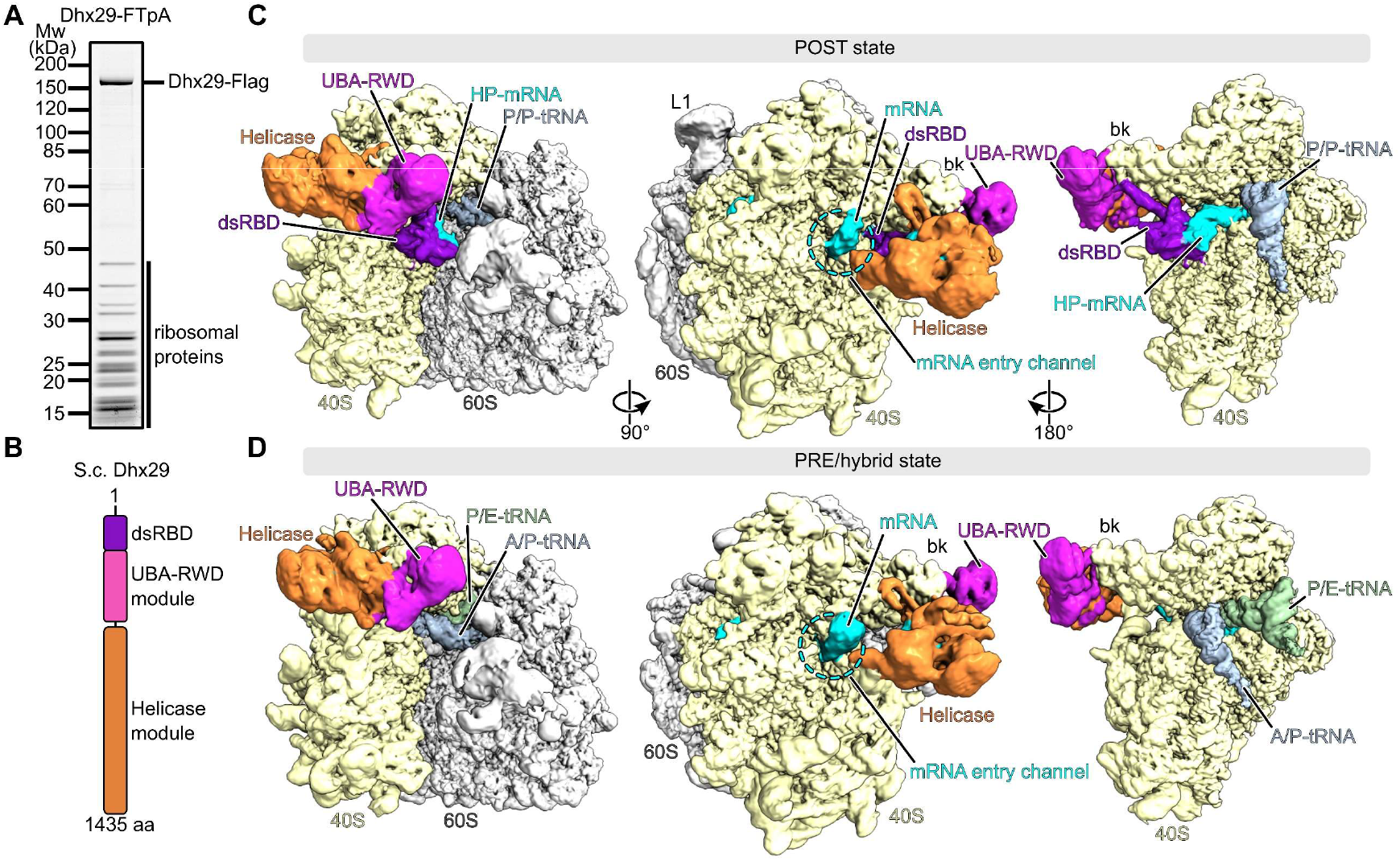
Cryo-EM structure of Dhx29 bound to translating 80S ribosomes. (**A**) SDS-PAGE gel showing the elution fraction of the Dhx29-FTpA affinity purification. Selected gel lane is cropped from gel shown in Fig. 2A. (**B**) Schematic of the main structural modules of Dhx29. (**C**, **D**), Cryo-EM densities of the Dhx29-80S complexes displayed from the front showing the intersubunit space (left), from the 40S solvent side focusing on the mRNA entry (middle), and from the 40S intersubunit side highlighting the tRNA binding sites (right, 60S density omitted for clarity) in POST state (**C**) and PRE/hybrid state (**D**). Local resolution filtered maps processed according to Appendix Fig. S1 are shown. bk = 40S beak, L1 = 60S L1 stalk, HP = hairpin.

Based on AF2 predictions (Fleming *et al*, 2025; Jumper *et al*, 2021) we assigned all domains of Dhx29 to the corresponding densities (Fig. EV1F,G). The C-terminal helicase module is positioned beneath the mRNA entry channel, packed against the 40S head (Fig. 1B,D; orange). The second module, composed of UBA-like and UBA-RWD domains (UBA-RWD module), binds to the 40S beak (Fig. 1B,D; magenta). For the POST state Dhx29-80S complexes, the N-terminal dsRBD reaches into the A-site (Fig. 1C; purple) and is in close contact with a strong density that corresponds to mRNA forming a hairpin structure (Fig. 1C; cyan). Since the human homolog of Dhx29 has been shown to resolve mRNA secondary structures in the 5’UTR of initiating mRNA on 43S/48S preinitiation complexes (Abaeva *et al*, 2011; Parsyan *et al*., 2009; Pisareva *et al*., 2008), Dhx29 may perform a similar function during elongation in yeast. Intriguingly, mRNA secondary structures are not only a challenge in translation initiation, but can also substantially slow down the elongation step (Wen *et al*, 2008). Structured mRNA elements in the ribosomal A-site may directly interfere with accommodation and efficient decoding by cognate tRNAs, as shown for single stranded, helix-forming poly-A stretches (Chandrasekaran *et al*, 2019; Tesina *et al*, 2020) and inhibitory di-codon combinations (Gamble *et al*, 2016; Tesina *et al*., 2020). Here, we hypothesize that the helicase activity of Dhx29 could function in resolving structured mRNA in the A- site after sensing the presence of an RNA hairpin via its N-terminal dsRBD domain, by applying pulling force on the 3’ end of the mRNA.

To further investigate the potential role of the helicase and the dsRBD modules, we affinity purified ribosomal complexes bound to a K633A P-loop/Walker A mutant version of Dhx29 that is deficient in either ATP binding, and/or E730Q/ Walker B mutant deficient in ATP hydrolysis (Saraste *et al*, 1990; Walker *et al*, 1982), as well as a ΔN (Δ221) mutant lacking the N-terminal dsRBD probing the A-site (Fig. 2A). We then analyzed them by cryo-EM, except for the Walker A+B double mutant. For an in-depth analysis of the contribution of each Dhx29 construct to individual classes, we combined all four (mutants and WT) datasets to perform extensive 3D classification with ∼1.6 million 80S particles (Appendix Fig. S2A). With this approach we confirmed the presence of the two main classes, the Dhx29- bound PRE/hybrid state and POST state 80S, from which high-resolution maps of 2.1 Å and 2.5 Å (Appendix Fig. S2H-K), respectively, were reconstructed. After local refinements on the Dhx29 helicase module (for the PRE/hybrid and POST state 80S) and on the 40S head and body regions (for the PRE/hybrid state 80S) (Appendix Fig. S2B-G) we obtained composite maps (Fig. 2B, bottom) that served as the basis for building and refining molecular models of Dhx29-PRE/hybrid-80S and Dhx29-POST-80S (Appendix Table S1).

**Fig. 2:**
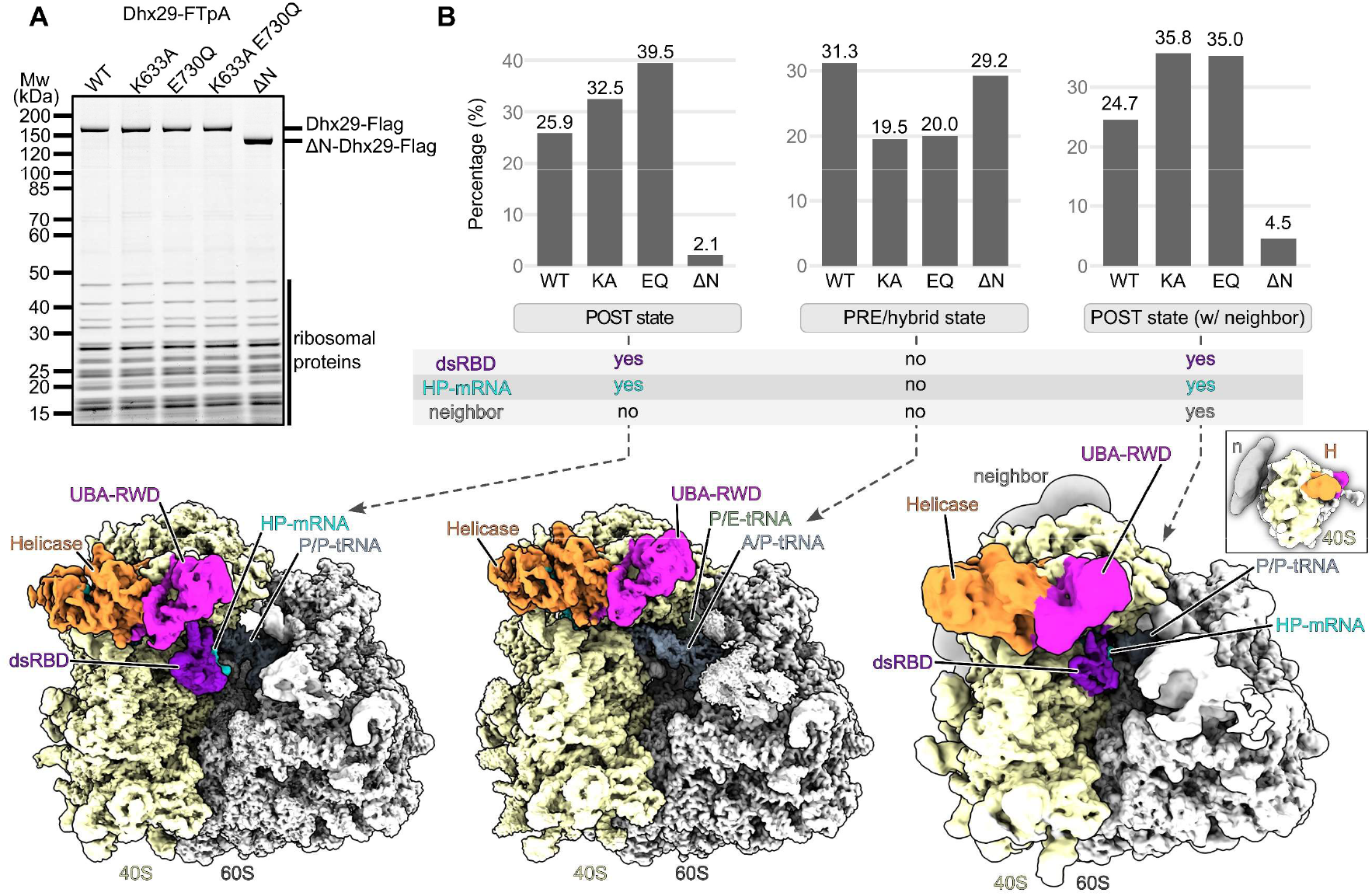
Cryo-EM analysis of 80S-bound Dhx29 mutants. (**A**) SDS-PAGE gel showing the elution fractions of the FTpA pulldowns for wild-type (WT) Dhx29, Walker mutants (K633A; Walker A, E730Q; Walker B) and an N-terminal deletion mutant (ΔN; Δ221). Lane for WT Dhx29 is also shown in Fig. 1A. (**B**) Single-particle cryo-EM analysis of Dhx29-80S complexes from the combined Dhx29 WT, K633A (KA), E730Q (EQ) and ΔN datasets, classified according to Appendix Fig. S2. Three classes - Dhx29-bound 80S in POST, PRE/hybrid states and Dhx29- bound POST state 80S involved in collisions - were analyzed for their relative contribution from the individual datasets. The diagrams indicate the percentage of particles from each individual dataset contributing to the three classes of the combined dataset. Shown below are composite cryo-EM maps of the Dhx29-80S complex in POST state and PRE/hybrid state and the local resolution filtered map of the POST-80S class showing density for a collided neighbor (n). Composite maps were obtained as described in the Methods section.

As observed in the wild-type sample (Fig. 1), the dsRBD of Dhx29 was present, bound to a mRNA hairpin, in the POST-80S complex, but absent in the PRE/hybrid-80S. Apart from those main classes, we observed classes of Dhx29-bound POST state 80S that showed density for a second ribosome at the mRNA exit site, indicating the presence of trailing ribosomes, likely representing ribosomal collisions (Appendix Fig. S2A). Interestingly, we observed neither idle ribosomes nor 80S bound to hibernation factors (Stm1/eEF2 and/or Lso2) (Li *et al*, 2024; Wells *et al*, 2020), indicating that Dhx29 binds specifically to active translating ribosomes.

To assess the relative differences in class distributions for WT and mutants, we matched the particles back to their parent datasets and compared their contribution to PRE/hybrid, POST or ribosome collision classes (POST-state 80S with neighbor density) (Fig. 2B and Appendix Fig. S3). Here, we observed a higher relative presence of Dhx29-bound ribosomes in POST state for both Walker A (K633A, short KA) and Walker B (E730Q, short EQ) mutants, when compared to the WT sample. We made a similar observation for the POST state 80S involved in collisions, which increase in number in both Walker mutants. The finding that these helicase deficient Dhx29 mutants accumulate stalled, colliding ribosomes is in agreement with our hypothesis that the Dhx29 ATPase is involved in resolving such stalls by employing its helicase activity to pull on the mRNA 3’ end. In the mutant lacking the N-terminal dsRBD, we observed opposite effects. Here, PRE/hybrid state 80S are significantly enriched, while POST states are essentially absent, indicating that, without this domain, Dhx29 is no longer able to specifically recognize hairpin-containing POST states and resolve collisions. Taken together, our data show that Dhx29 engages translating 80S ribosomes, which contain an mRNA, independent of an intact ATPase engine. The N-terminal dsRBD provides higher overall affinity and specificity for ribosomes in the POST state with a hairpin-like structured mRNA present in the A-site. Both the accumulation of ribosomes with an RNA hairpin in the A-site and the accumulation of collided ribosomes with helicase-deficient Dhx29 mutants strongly support the hypothesis of Dhx29 functioning in resolving A-site mRNA structures for efficient progression of elongation and to avoid stress-triggering collisions.

### Dhx29 engages the ribosome via its UBA-RWD module and OB domain insertions

In the molecular model of both PRE/hybrid (Fig. 3A) and POST state translating Dhx29-80S complexes, the helicase module is located between the 40S body and head, approximately 60 Å above the mRNA entry site. We observed density for mRNA from the 5’ mRNA exit site, through the mRNA channel, to the 3’ entry site and all the way into the helicase core when low-pass filtering the isolated mRNA density (Fig. 3B). In more detail, we could model twenty nucleotides of the mRNA located within the mRNA channel of the 40S, and ten mRNA bases could be clearly traced and built within the Dhx29 helicase module of the 80S PRE/hybrid state (Fig. 3B). Both parts are connected via a more flexible, bulky mRNA density. In order to span the stretch between the 40S mRNA entry and the helicase entry site, at least 6-12 nucleotides would be required (straight path or following the shape of the bulky density, respectively), totaling about 36-42 nucleotides in the Dhx29-80S hybrid state complex. Notably, at lower contour levels, we also observe further extra density for the 3’ region of the mRNA that exits the helicase and appears to fold back onto the top of the helicase module on a positively charged groove/surface, similar to the *E. coli* RNA helicase HrpA when bound to collided disomes (Campbell *et al*., 2025) (Appendix Fig. S4A,B).

**Fig. 3:**
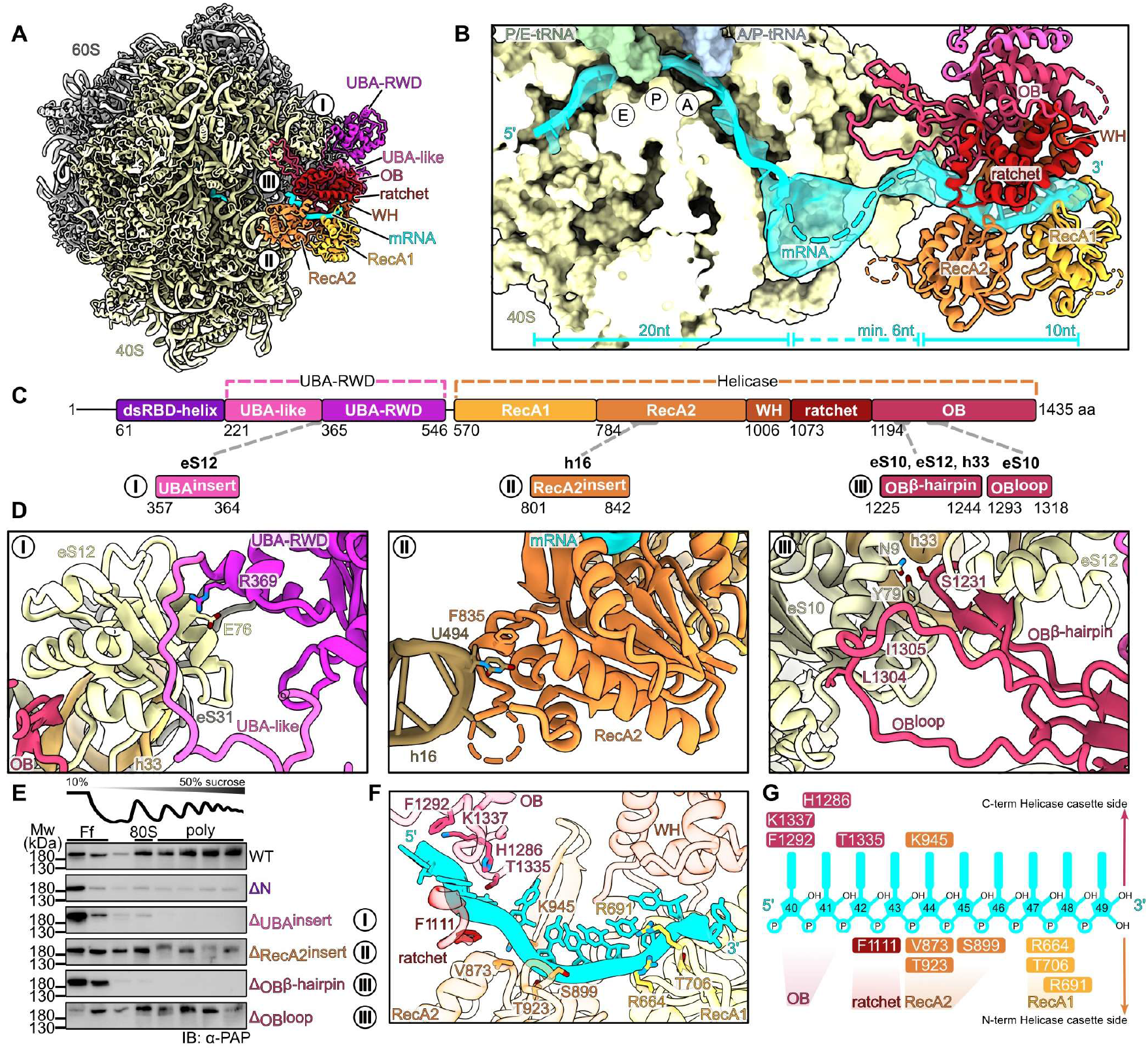
Molecular model of Dhx29 helicase and UBA-RWD module contacts with PRE/hybrid state 80S. (**A**) Cryo-EM based molecular model of the Dhx29-80S complex in PRE/hybrid state, shown as a top view. Regions for zoom views shown in (**D**) are indicated (I-III). (**B**) Zoom highlighting the mRNA path from the 5’ mRNA exit site beyond the 3’ mRNA entry site of the 40S into the Dhx29 helicase core. Isolated low-pass filtered density for mRNA is shown transparent, the model for the 40S is shown in surface representation and the helicase module as ribbons. (**C**) Diagram of structural elements/domains of Dhx29. Regions involved in ribosome binding are indicated (I-III). (**D**) Zoom on the UBA-RWD module contacting eS12 (I), RecA2 insert bound to the tip of h16 (II) and the two OB domain contact sites (OB-β-hairpin and OB-loop) with the groove formed by eS10, eS12 and h33 (III). (**E**) Western blots showing the distribution of Dhx29 ribosome-binding mutants across sucrose gradient fractions. Ff = free fraction; poly = polysomes. (**F**) Close-up of the Dhx29–80S PRE/hybrid state model showing mRNA engagement by conserved residues within the helicase module. (**G**) Schematic representation of the interactions shown in (**F**).

The Dhx29 helicase module employs similar interaction sites on the 40S (Appendix Fig. S5A) as other ribosome-bound helicases, including human DHX29 (Cui *et al*, 2025; Hia *et al*., 2026) (Appendix Fig. S5B,C), yeast Ski2 (Schmidt *et al*., 2016) (Appendix Fig. S5D) and Slh1 (Best *et al*., 2023) (Appendix Fig. S5E), as well as bacterial HrpA (Campbell *et al*., 2025) (Appendix Fig. S5F). These sites include the 18S rRNA helix 16 (h16) at the 40S body, as well as the 40S head. Three anchors place the Dhx29 helicase module on the 40S (Fig. 3C,D). The 40S body anchor is formed by an insertion within the RecA2-domain (residues 801-842) which, in line with previous crosslinking data (Fromont-Racine *et al*., 2024), engages the 18S rRNA helix h16. It wraps around the tip and stem of h16 (visible only at low contour levels) and is positioned by a short α-helix (residues K834-T837) binding to the tip of h16 (formed by U493, U494 and U495) via F835 (Fig. 3D, II and Fig. EV2A). Two prominent protrusions emerge from the OB domain and establish contacts with the 40S head and beak (Fig. 3D, III and Appendix Fig. S4C-E). The first protrusion forms a β-hairpin (residues 1225-1244; OB-β-hairpin) that reaches into a pocket formed by ribosomal proteins eS12, eS10 and 18S rRNA helix h33, with clear contacts established by S1231 with Y79 and N9 of eS10. The second protrusion forms an unstructured loop (residues 1293-1318; OB-loop) which binds mainly via the hydrophobic residues L1304 and I1305 contacting eS10. The helicase module of Dhx29 is connected via an unstructured linker (residues 547-569) to the combined UBA- RWD domain (residues 365-546) preceded by the UBA-like domain (residues 221-364) that partially resembles the N-terminal domain of the human DHX29, which packs against the OB fold of the helicase module (Fig. EV1A,B,D). This whole structural block (the UBA-RWD module) is packed against eS12 and is close to the 40S beak protein eS31. The primary contacts are made by a loop region of the UBA-like domain that connects to the UBA-RWD domain (residues 357-364) establishing mostly backbone interactions with eS12 (Appendix Fig. S4F-H). Additionally, a weak specific contact is established between R369 and E76 of eS12. (Fig. 3D, I).

A direct comparison of the engagement of the 40S by the yeast Dhx29 and the human DHX29 helicase modules (Hia *et al*., 2026) reveals distinct differences (Fig. EV2). While yeast Dhx29 only contacts h16 via the RecA2 insert (Fig.3D, II and Fig. EV2A), human DHX29 interactions with h16 are more prominent and are established by the functionally important RecA2 β-hairpin, as well as by the WH and OB domains (Fig. EV2D). The 40S head contacts of yeast Dhx29 are formed via the two loop inserts within the OB domain and the UBA-RWD module (Fig. EV2B). However, the UBA-RWD domain, as well as the OB-β-hairpin insert, are missing in human DHX29 (see also Fig. EV1C,E). As a result, contacts with the 40S head are limited to one shorter loop of the OB domain and the N-terminal helix of the UBA-like domain contacting eS12 on the 40S beak (Fig. EV2E).

To test how the described contacts contribute to the interaction of Dhx29 with the ribosome, we individually deleted the contact sites with h16 (RecA2 insert, Δ803-828-GSGSG), the eS10/eS12/h33 pocket (OB-β-hairpin, Δ1225-1237-GSGS), eS10 (OB-loop, Δ1296-1314-GSGSGS), and the eS12/eS31 contact site between the UBA-like and UBA domains (UBA-insert, Δ357-364-GSGSGSGS). In addition, we tested the N-terminal deletion mutant (ΔN) lacking the first 211 residues. All Dhx29 constructs were expressed from monocopy plasmids under the control of the endogenous *DHX29* promoter and fused to a C-terminal FTpA-tag in *dhx29*Δ cells. Cell lysates were applied to sucrose density gradients and fractions were analyzed by Western Blotting against the protein A tag (Fig. 3E). This experiment showed that Dhx29 mutants with deletions in either the UBA-insert (eS12/eS31-contact) or in the OB- β-hairpin almost completely lose their ability to bind to polysomes compared to the wild-type, while the RecA2 insert and the OB-loop mutants showed no or only minor effects in polysome binding. Interestingly, contrary to the OB-β-hairpin and UBA-insert mutants, the ΔN mutant was still able to interact with ribosomes, although at strongly reduced levels. This suggests that the ΔN mutant may have lost its specificity but still displays some remaining affinity for ribosomes. This is in line with the cryo-EM analysis of the ΔN mutant 80S complex showing a severely decreased association with POST state 80S and increased enrichment of the PRE/hybrid state ribosomes (Fig. 2B).

### Dhx29 mRNA binding is conserved with DEAH box 3’-5’ helicases

In all our structures, irrespective of PRE or POST state, we find the helicase module engaged with mRNA, bound between the cleft formed by the two RecA domains on one side and the WH/Ratchet/OB domains on the other side. The conformation resembles the one of the RNA-bound DEAH-box spliceosomal helicase Prp22 from *Chaetomium thermophilum* (Hamann *et al*, 2019), representing an open conformation with respect to the two RecA domains, with no nucleotide density between the RecA domains (Fig. EV3A, left panels). Conversely, structures of Prp43-ADP-BeF₃ that are arrested in closed RecA domain conformation, with or without RNA (Hamann *et al*., 2019; Tauchert *et al*, 2017), are not compatible with our cryo-EM density (Fig. EV3A, right panels).

Within the Dhx29 helicase module, mRNA binding is mediated by several residues in the RecA domains on the RNA backbone-facing side of the cleft, and by the WH, ratchet and OB domains on the opposite, base-facing side (Fig. 3F,G). The comparison with the X-ray structure of Prp22 bound to single-stranded RNA (Fig. EV3B) shows that most of the RNA-interacting residues are conserved in the Dhx29 RecA domain, including the residues that contact the mRNA backbone (i.e., R664 in RecA1; S899, N924 and K945 in RecA2) as well as the residues that contact the 5’-bases of the mRNA in the OB and ratchet domains (via H1286, F1292, T1335 and K1337 in OB and F1111 in the ratchet domain). The path of the 3’ end of the mRNA deviates towards the Dhx29 RecA2 domain when compared to Prp22. Overall, Dhx29 engages the mRNA in a manner similar to other DEAH box helicases, which translocate in the 3’-5’ direction with a step-size of one nucleotide per hydrolyzed ATP (Boneberg *et al*, 2019; Hamann *et al*., 2019).

### The Dhx29 dsRBD domain senses mRNA hairpins in the A-site

The dsRBD containing N-terminal region of Dhx29 was not visible in any of the PRE state ribosomal classes, indicating that it is not stably bound to the ribosome and is rather flexible in this state. However, the dsRBD becomes stably positioned in POST state ribosomes that contain a problematic, structured hairpin-like mRNA in the A-site. In this state the dsRBD is located in the translation factor binding site of the 40S (Fig. 4A,B) and directly binds to the strong and compact density that corresponds to the mRNA hairpin in the A-site (Fig. 4B). The Dhx29/DHX29 (yeast/ human) dsRBD consists of the conserved α1-β1-β2-β3-α2 fold found in all dsRBDs (Masliah *et al*, 2013) (Fig. 4C). It is preceded by an N-terminal α-helix (α0) and followed by three α-helices (α3-α4-α5), which mediate contacts with the 40S subunit. The long α5 helix links the dsRBD to the UBA-RWD module. Overall, the dsRBD interacts with the 40S through contacts with uS12 and the N-terminus of eS30. This interaction is similar to that observed for the counterpart in human DHX29 (Hia *et al*., 2026) (Fig. EV2C,F). However, the two dsRBDs differ in the conformation of their long C-terminal α-helix that bends towards h16 in case of the human DHX29, while it stretches out towards h33 and the 40S beak for yeast Dhx29 (Fig. EV2C,F). More specifically, the yeast Dhx29 dsRBD interacts with the 40S through T55 and S57 (to uS12 D97), T59 (to uS12 K56) and N152 (to the backbone of eS30 residues K15, S16, T18) (Fig. 4C, I). The long α5 helix (residues 191-220) is partially visible and stretches along h33, forming contacts to the phosphate-backbone of A1226 (by Q215), U1225 and A1224 (by E208). The flipped-out G1228 is contacted by the backbone of Q217 and K218, as well as T67 of eS12 (Fig. 4C, II). The positioning of the Dhx29 dsRBD exposes a positively charged surface, which is mainly formed by α1 and the flanking regions towards the A-site decoding center (Fig. 4C, III and Fig.4D). This surface is juxtaposed to the mRNA hairpin in the A-site. Based on our consensus reconstruction, we replaced the A-site codon by a 26 nt long mRNA structure that forms a 22 nt hairpin capped by a GAAAC tetraloop and modeled it into this density. Its structure resembles mRNA HPs previously observed in context of translational bypassing (Phage T4 *gene60*) (Agirrezabala *et al*, 2017) and programmed frameshifting (HIV-FSS) (Bao *et al*, 2020) in bacteria, as well as an mRNA loop formed by an artificial CGA-CCG mRNA di-codon sequence in yeast POST state ribosomes (Tesina *et al*., 2020) (Fig. EV4A).

**Fig. 4:**
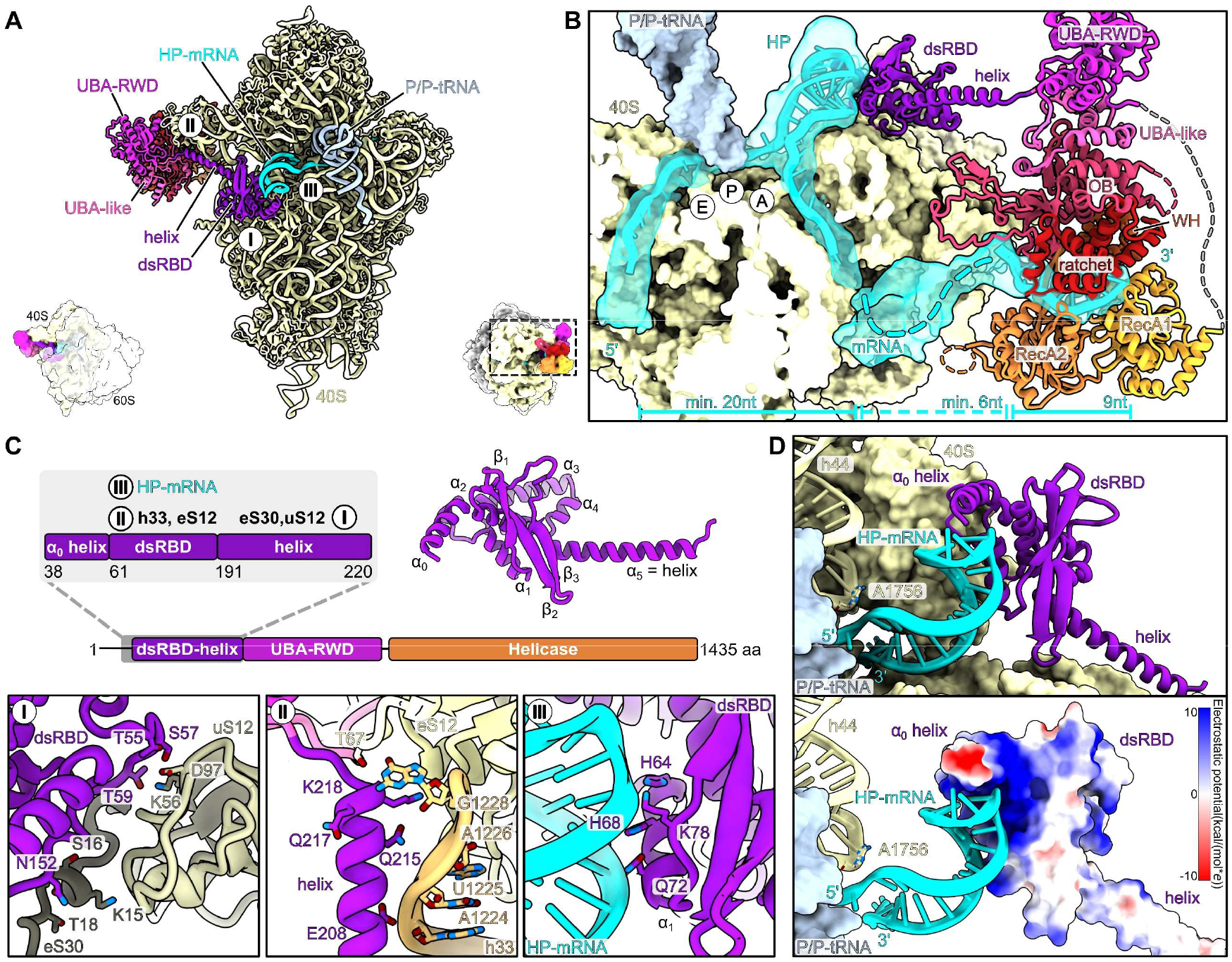
Molecular model of contacts between the Dhx29 N-terminal module and the POST state 80S. (**A**) Cryo-EM-based molecular model of the Dhx29-80S complex in POST state, shown in side-view. A thumbnail indicates the orientation. Regions for zoom views shown in (**C**) are indicated. (**B**) Zoom on the mRNA channel as shown in Fig. 3B. The mRNA forms a hairpin (HP) in the decoding center and binds to the Dhx29 dsRBD. Isolated low-pass filtered density for mRNA is shown transparent, the model for the 40S is shown in surface representation and the helicase module as ribbons. (**C**) Top: domain organization of Dhx29 with focus on the N-terminal dsRBD (left) and molecular model (right) with hallmark secondary structure elements labelled. Bottom: zoom views highlighting the interaction of dsRBD domain with 40S proteins uS12 and eS30 (I), the (linker) helix contacting eS12 and h33 (II) and the Dhx29 dsRBD with the mRNA HP (III). (**D**) Zoom on the contact interface between the minor groove of the HP-mRNA and positively charged residues of the dsRBD domain (III). Top: ribbon model; bottom: dsRBD surface colored according to the electrostatic potential.

Further sub-classification with focus on the HP region revealed subsets of even longer HPs, indicating that the consensus reconstruction likely contains a mixture of mRNA structures that vary in sequence and length (Fig. EV4B). However, the particle numbers in these sub-datasets were too low for high- resolution structure determination and detailed molecular interpretation. As observed in other RNA- bound dsRBD structures, like the *Xenopus laevis* dsRBD-A (Masliah *et al*., 2013; Ryter & Schultz, 1998), the main contact with the A-helix of the hairpin is established by α1, which has been shown to mediate sequence-unspecific, shape-based recognition of RNA A-helices by binding into the minor groove (Fig. EV4C). Within the Dhx29 dsRBD α1 helix side chain densities are resolved for H64, H68 and Q72 and the loop region between α1 and β1 that contributes to the HP interaction via K78 which is contacting the phosphate backbone (Fig. 4C,III). Additional contacts are formed by the N-terminal α0, which is located at the tip of the HP (Fig. 4D). However, this helix appears more flexible and likely adapts to the size and shape of the HP.

Taken together, in the event that an mRNA hairpin folds within the A-site, the dsRBD of Dhx29 recognizes its presence, while a tRNA in the A site (or in the A/P site) would be out of reach (Fig. EV4D). This interaction may therefore serve as a sensor for structured mRNA in the A-site, eventually triggering the Dhx29 helicase domain to pull the mRNA in 5’-3’-direction, resulting in unwinding of the hairpin. This would be consistent with human DHX29, in which deletion of the dsRBD suppresses the NTPase stimulation by the 40S subunit (Sweeney *et al*, 2021).

### Selective ribosome profiling of Dhx29-bound ribosomal complexes

To learn more about the endogenous mRNA substrates of Dhx29, we performed selective ribo-seq using the *DHX29*-FTpA strain and compared it to ribo-seq of a *DHX29* deletion (*dhx29*Δ) strain. Comparing the metagene profiles of selective Dhx29 ribo-seq with the global input and the *dhx29*Δ ribo-seq, the overall profiles were largely similar, without pronounced positional preferences for Dhx29 across the CDS (coding sequence) (Fig. 5A). In the *dhx29*Δ strain, however, the stop-codon peak was markedly reduced, while a peak approximately 30 nt upstream of the stop codon was increased relative to the WT global input control. Notably, both positions also showed the strongest Dhx29 enrichment in the selective ribo-seq metagene profile. Comparing footprint length distributions across the three conditions, selective Dhx29 ribo-seq yielded substantially longer footprints than either the WT global input or the *dhx29*Δ samples (Fig. 5B), consistent with extended protected mRNA due to secondary structure formation in the A-site of the ribosome or protection by the helicase itself. Mapping footprint length back onto our structural analysis, the observed length of ∼36 nt and longer agrees with a cleavage site immediately downstream of or within the stem loop positioned in the A- site of Dhx29-engaged ribosomes (Fig. 5C). Differential enrichment analysis (DESeq2) identified 505 transcripts passing our cutoffs of log2FC > 1 and −log10(Padj) > 10 (Fig. EV5A). GO term enrichment revealed a modest but significant bias towards transcripts encoding ribosomal proteins among Dhx29- engaged ribosomes (Fig. EV5B). Seeking additional features that might explain preferential Dhx29 recruitment, we related enrichment to CDS length and found that shorter CDSs had a higher propensity to recruit Dhx29 (Fig. EV5C). A recent study in human cells proposed that DHX29 acts as a codon optimality sensor, being preferentially recruited to codons carrying A or U at the wobble position (AU3) (Hia *et al*., 2026). We therefore asked whether yeast Dhx29 behaves similarly, however, AU3 (inverse GC3) content correlated only weakly with Dhx29 enrichment (Fig. EV5D), whereas the tRNA adaptation index (tAI) as a readout of codon optimality showed a stronger correlation (Fig. EV5D,E). Low GC1 content, in contrast, was still a weak yet comparably stronger predictor of Dhx29 recruitment (Fig. EV5D), suggesting that amino acid identity rather than synonymous codon choice may contribute to recruitment. Consistently, the hydrophobicity of the encoded amino acids predicted Dhx29 recruitment similarly well (Fig. EV5D).

**Fig. 5:**
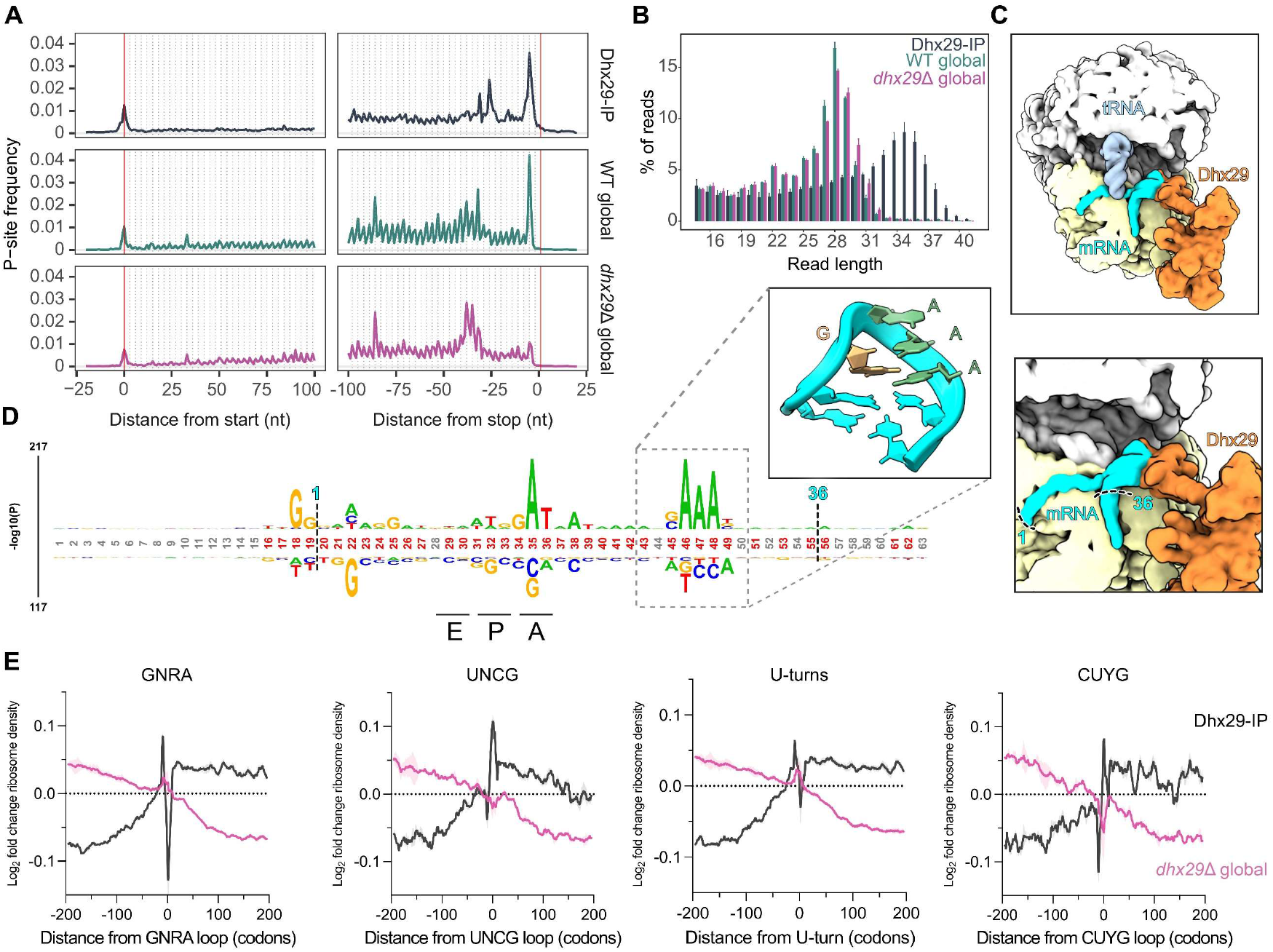
Selective ribosome profiling of Dhx29 complexes and a *dhx29*Δ strain links Dhx29 to structured mRNA elements. (**A**) Metagene profiles of selective Dhx29 ribosome profiling and of global ribosome profiling from wild-type (WT) and *dhx29*Δ cells. (**B**) Read-length distributions of ribosome-protected fragments for the indicated samples. (**C**) Structural overview of the mRNA path in the model (top, cyan) and the corresponding median footprint length mapped onto it (bottom). (**D**) Sequence logos around stall sites in *dhx29*Δ relative to WT, revealing a GNRA tetraloop motif downstream of the A-site. **(E**) Structured loop elements (GNRA, UNCG, U-turns, CUYG) are associated with increased ribosome density upstream of the motif in the absence of Dhx29, consistent with Dhx29 resolving these elements during elongation.

We next addressed the translational consequences of *DHX29* deletion. Differential expression analysis revealed minimal global changes in translation, with the exception of Dhx29 itself, confirming our deletion strain (Fig. EV5F). Based on our structural work, we hypothesized that the helicase activity of Dhx29 resolves secondary structures occluding the A-site and thereby promotes ribosome progression. We therefore determined at which positions ribosomes pause in the absence of Dhx29. Examining the sequence context around these pause sites, we found a strong bias for a [G/C]AAA[G/C] motif downstream of the E-, P- and A-site codons (Fig. 5D), reminiscent of GNRA tetraloops (Bujnicki & Baulin, 2025). We next asked whether such structures recruit Dhx29 and whether Dhx29 is able to resolve them. GNRA loops were indeed enriched in Dhx29 footprints (Fig. 5E, grey lines), and loss of Dhx29 increased ribosome occupancy immediately upstream of the GNRA loop (Fig. 5E, pink line). Extending this analysis to other well-characterized RNA loops, UNCG loops, U-turns and CUYG (Y = C or T) loops (Bottaro & Lindorff-Larsen, 2017) all showed a comparable pattern of Dhx29 enrichment and upstream ribosome accumulation upon *DHX29* deletion (Fig. 5E). This points again towards a role of Dhx29 in resolving structured mRNAs containing relatively stable loops (as parts of hairpins) that otherwise would slow ribosome progression. Dhx29 recruitment followed codons at the A-site with a lower tRNA adaptation index (R = -0.36, P= 0.01), consistent with a recruitment to slowly decoding ribosomes. However, ribosomes were not more prone to stall at these codons in a *dhx29*Δ mutant strain (R = 0.07, P = 0.6) (Fig. EV5G). Similarly, we observed only a weak correlation at E-site codons between recruitment and stalling if Dhx29 is absent (R = 0.30, P = 0.02), uncoupling recruitment and function at A- and E-site codons. In contrast, P-site codons showed a strong correlation between recruitment and stalling (R = 0.62, P < 1 × 10⁻¹⁶), with a bias for pyrimidines at position 1. Together, these data suggest that Dhx29 recruitment depends not only on RNA secondary structures downstream of the decoding center, but also on the identity of the codon occupying the P-site.

## Discussion

This work presents a structural and functional characterization of Dhx29 (Ylr419w), the yeast homolog of the ATP-dependent 3’-5’ RNA helicase DHX29 (Fromont-Racine *et al*., 2024). Cryo-EM analyses of native Dhx29-ribosome complexes suggest a role of Dhx29 in maintaining translation progression by unwinding stable mRNA structures (hairpins and/or stem-loops) that form in the A-site (Fig. 6). Selective ribo-seq confirms this finding and suggests that Dhx29 is enriched on ribosomes engaged upstream of tetraloops. Yet, our data hint at Dhx29 functions beyond resolving stable mRNA secondary structures. Dhx29 recruitment shows a weak but significant correlation with the tRNA adaptation index (tAI), indicating that Dhx29 senses slowed ribosomes. Globally, we find Dhx29 enriched on transcripts with rather shorter CDS, such as transcripts encoding ribosomal proteins.

**Fig. 6:**
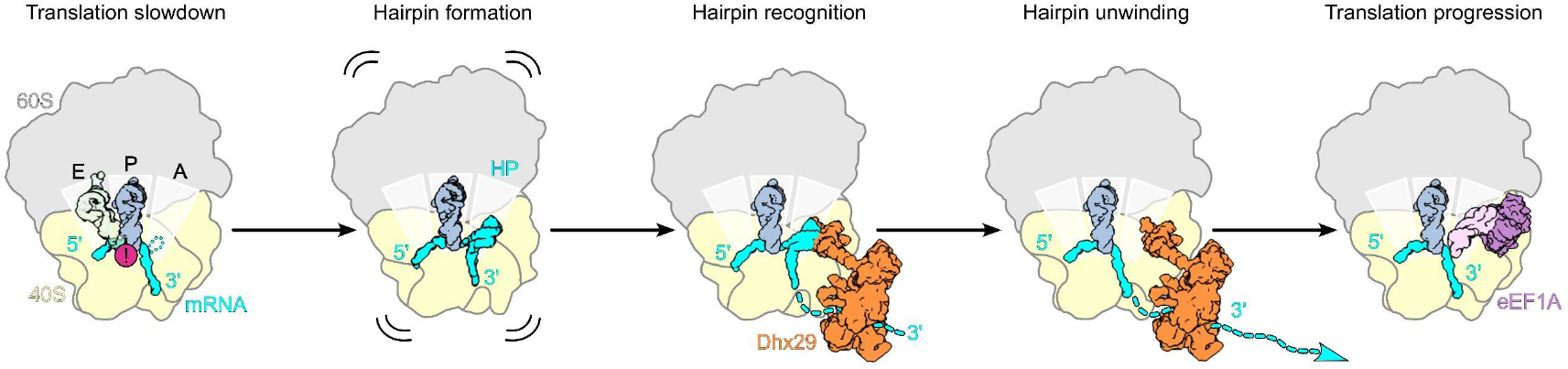
Schematic representation of mRNA hairpin recognition and unwinding by Dhx29 on the ribosome. Hairpins (HPs) can form spontaneously in the ribosomal A-site of slowed or stalled ribosomes (indicated by the exclamation mark) based on tetraloop-prone mRNA sequences. Dhx29 binds translating ribosomes via 3’ mRNA overhangs and positions its dsRBD in a conformation that senses the presence of such HP. The helicase engine of Dhx29 pulls the mRNA in 5’-to-3’ direction, thus unwinding the HP. Following clearance of the A-site, translation can proceed to the next decoding step.

When considering our ribo-seq data together with our structural analysis, it is important to note that the majority of Dhx29-bound 80S ribosomes are found in the PRE/hybrid state where we do not observe any mRNA-HPs, nor dsRBD engaged with these ribosomes. This, in turn, means that general effects detected in the profiling data likely reflect the nature of the hybrid state ribosomes that have no general decoding problem (A/P site tRNA is accommodated), but may be slow in translocation. Thus, we hypothesize that Dhx29 can engage ribosomes for global monitoring of translation, an interaction that may occur independently of the presence of an mRNA hairpin in the A-site and independent of the dsRBD-containing N-terminal domain. In addition, ribosome and mRNA binding are not impaired by the Dhx29 Walker A and B mutants, indicating that engagement is unrelated to its helicase activity.

In ATPase mutants, however, POST states with HP density in the A-site accumulated, indicating that ATP-hydrolysis is required to resolve such HP structures. In case the N-terminal dsRBD is lacking, the PRE/hybrid states are enriched, suggesting that this deletion mutant lost its specificity to interact with 80S harboring HP structures in the A-site, a particle species that was essentially absent in the Dhx29- ΔN-80S dataset.

Our results indicate that yeast Dhx29 has a similar function as shown for human DHX29 during translation initiation. DHX29 was first described to unwind structured mRNAs during the scanning process of translation initiation as a component of the 48S initiation complex (Pisareva *et al*., 2008). Also in the 48S complex, DHX29 sits at the mRNA entry site, positioning its dsRBD in the A-site, where it could monitor the formation of HP structures (Cui *et al*., 2025; des Georges *et al*, 2015; Hashem *et al*., 2013). A recent study proposed that - similar to our observations - the DHX29 dsRBD might bind to HPs in the A-site of the scanning 48S complex, outcompeting eIF1A, which leads to closure of the mRNA latch and stops scanning. After pulling on the mRNA and resolving problematic secondary structures, the helicase would dissociate, enabling eIF1A to bind and scanning to resume (Cui *et al*., 2025). In contrast, the recently suggested function of human DHX29 on translating 80S ribosomes for recognition of non-optimal codons (and subsequent mRNA decay) (Hia *et al*., 2026) seems not to be shared with yeast Dhx29. Our selective ribo-seq data did not reveal any enrichment of non-optimal codons in the A-site and it is unclear how the hairpin-binding dsRBD of Dhx29 would recognize the nature of the A-site codon. It is therefore unlikely that the yeast Dhx29 plays a role in codon optimality sensing. An additional difference between human and yeast Dhx29 is the absence of the UBA-RWD domain in human DHX29. RWD domains are generally known to mediate the interactions of proteins with their respective binding partners, as in the case of the co-translational stress sensors Gcn2 and Gcn1 (Kubota *et al*, 2000; Sattlegger & Hinnebusch, 2000). Therefore, it is reasonable to speculate that yeast Dhx29 may additionally modulate the integrated stress response (ISR) when bound to the ribosome via direct competition with the Gcn2-RWD domain. Nevertheless, under the applied growth conditions in rich media and without stress, we did not observe Gcn1 or other ISR-related factors co- enriching with the native Dhx29 purifications.

Taken together, our results lead to a model that Dhx29 generally associates with translating 80S ribosomes via its helicase and UBA-RWD modules to globally monitor translation. In case of ribosome slowdown, mRNA in the A-site can rearrange, as shown for poly-(A) stretches or in case of inhibitory di-codons (Tesina *et al*., 2020) and eventually form HP structures occupying the A-site. The Dhx29 dsRBD senses such HP structures that are subsequently unwound by the helicase module, pulling at the mRNA in 3’-5’ manner, opposite to the direction of translation. This would resolve the mRNA HP in the A-site, lining up the codon triplet for recognition by the next ternary aminoacyl-tRNA-eEF1A- GTP complex, thus resuming translation elongation and maintaining translation progression.

## Methods

### Plasmids

Plasmids used in this study are listed in Appendix Table S2. DNA cloning was performed with PCR amplification using gene-specific primers and Platinum SuperFi II polymerase (Thermo Fisher). Fragments corresponding to the *DHX29* promoter and CDS (wild-type sequence or carrying primer introduced mutations) were digested using restriction enzymes EcoRI, NdeI and BamHI (NEB), then ligated into a YCplac111 backbone (Rizzardi *et al*, 2012) containing a C-terminal 5xGA-FTpA tag using T4 DNA ligase (NEB). All cloned DNA sequences were verified and confirmed by sequencing.

### Yeast strains

The *Saccharomyces cerevisiae* strains used in this study were derived from W303-1a (Thomas & Rothstein, 1989). The *DHX29* deletion (*dhx29*Δ) and the *DHX29*-FTpA strains were constructed using homologous recombination of PCR-amplified cassettes and selected using the *natNT2* selection marker (Janke *et al*, 2004; Longtine *et al*, 1998). Strains were confirmed by fragment PCR amplification and sequencing. Plasmids containing WT, mutant, or truncated Dhx29 variants fused to a C-terminal FTpA tag (Flag - TEV protease cleavage site - protein A) were transformed into the *dhx29*Δ background (see Appendix Table S2 and S3) and positive colonies were selected by growth on SDC-Leu plates.

### Affinity purification of Dhx29-ribosomal complexes

The Dhx29 samples used for the single-particle cryo-EM analysis were purified from *dhx29*Δ yeast cells that were transformed with the respective Dhx29-FTpA wild-type and mutant constructs (see Appendix Table S2). The cells were grown at 30 °C with constant shaking at 125 rpm in SDC-Leu medium until an OD_600_ of ∼2 was reached. The cells were harvested by centrifugation, diluted in fresh YPD medium, and grown for an additional six hours at 30 °C until an OD_600_ of 2.0–2.4 was reached. The yeast cell pellets were frozen in liquid nitrogen as drops and then milled with a SPEX 6970EFM Freezer/Mill. The cell powder was resuspended in purification buffer (50 mM Tris-HCl pH 7.5, 60 mM NaCl, 40 mM KCl, 5 mM MgCl₂, 1 mM DTT) supplemented with 5% glycerol, 0.1% IGEPAL CA-630 (Sigma-Aldrich) and homemade protease inhibitor cocktail. The lysate was pre-clarified by two centrifugation steps: one at 3,095 x g for 10 min at 4 °C, and one at 36,500 x g for 25 min at 4 °C. For the first affinity purification step, IgG Sepharose 6 Fast Flow resin (Cytiva) was added to the cleared lysate, and binding was performed at 4 °C for 90 minutes under rotation. The resin was then collected and washed once in batch, followed by a wash step using a 20 mL Econo-Pac chromatography column (Bio-Rad). The IgG beads were collected, and the samples were eluted by incubating with homemade TEV protease at 16°C for 90 min. For the second affinity purification step, the TEV eluate was incubated with pre-equilibrated anti-FLAG M2 affinity gel (Sigma-Aldrich) at 4 °C for 90 min under rotation. The resin was collected and washed once in purification buffer supplemented with 0.01% IGEPAL CA-630 (Sigma-Aldrich), followed by a second wash with purification buffer supplemented with 0.05% octaethylene glycol monododecyl ether (Nikkol). The resin was then transferred to a 1-mL Mobicol gravity-flow column and washed with 5 mL of buffer containing 0.05% octaethylene glycol monododecyl ether (Nikkol). Samples were eluted by adding 250 μg/mL of 3xFLAG peptide and incubating for 60 minutes at 4°C.

### Polysome analysis

For *in vivo* binding assays, yeast cells were grown in SDC-Leu medium with 2% glucose at 30°C and 125 rpm constant shaking, and harvested at log phase (OD_600_ ∼ 0.8). Prior to harvesting, cells were quickly cooled in an ice bucket and pre-treated with 100 μg/mL cycloheximide (Sigma Aldrich, C7698) for 10 min before centrifugation. Cell pellets were washed, then resuspended in lysis buffer (50 mM Tris-HCl pH 7.5, 150 mM KCl, 12 mM MgCl_2_, 1% Triton-X, 1 mM DTT, 1 mM PMSF, 100 μg/mL cycloheximide, supplemented with homemade complete protease inhibitor and RNase inhibitor). Cells were mechanically lysed using 0.5 mm glass beads, in 10 cycles of 30 s vortexing followed by 30 s cooling on ice. The clarified lysate was collected upon a short spin and a centrifugation at 20,817 x g, 4°C for 15 min. Absorbance at 260 nm (A_260_) was measured and an equivalent of 8 A_260_ units (320 μg RNA) was loaded onto a 10-50% sucrose gradient (prepared in 50 mM Tris-HCl pH 7.5, 100 mM KCl, 12 mM MgCl_2_, 1mM DTT) for each sample. Sucrose gradients were subject to centrifugation at 243,792 x g and 4°C for 150 min in a SW 40 Ti rotor (Beckmann Coulter). Then, sucrose gradients were fractionated using a gradient station coupled to an automated fractionator (BioComp). Proteins from collected fractions (free, ribosomal subunits, monosomes and polysomes) were precipitated by TCA (10% final concentration) and washed twice with acetone before drying. Pellets were resuspended in SDS-PAGE loading buffer (50 mM Tris-HCl pH 6.8, 100 mM DTT, 8% SDS, 0.1% bromphenol blue, 10% glycerol). Resulting samples were loaded on 10% NuPAGE gels followed by Western Blotting.

### Electrophoresis and Western blotting

Protein samples from affinity purifications were separated by SDS-PAGE on 4-12% NuPAGE gels (Thermo Fisher) and gels were stained with Coomassie (Blauer Jonas, GRP). TCA precipitated samples from sucrose gradients were separated by SDS-PAGE on 10% NuPAGE gels and transferred to 0.45 μm PVDF membranes (Immobilon-P, Millipore). Membranes were then blocked using 2% skim milk in 1x PBS solution and incubated with peroxidase-anti-peroxidase complex antibody (PAP-HRP 1:3000, Sigma P1291) for an hour at room temperature. After washing two times with 1x PBS-T and once with 1x PBS, the membranes were incubated with ECL substrate (Thermo Fischer) and chemiluminescence was detected using LAS4000 mini (GE Healthcare).

### Cryo-EM sample preparation and data collection

Freshly prepared samples (∼ 3.0 A_260_/mL) were applied to glow-discharged R3/3 holey copper grids coated with a 3 nm continuous carbon support film (Quantifoil) using a Vitrobot Mark IV (FEI Company) operated at 4 °C and 90% humidity. The samples were incubated for 45 s on grids, then excess sample was blotted for 3 s, followed by plunging in liquid ethane. Cryo-EM data was collected on a 300 kV Titan Krios G3 (FEI Company) equipped with a Falcon 4i direct electron detector and a Selectris X energy filter (Thermo Fischer) under low-dose conditions (40 frames per movie, total dose of 40 e^−^/Å^2^). Micrographs were collected in EPU (Thermo Fischer) at 165,000 magnification, resulting in a pixel size of 0.727 Å. The defocus used ranged between 0.5 to 3.5 μm. All frames were gain-corrected, aligned and subsequently summed using MotionCor2 (Zheng *et al*, 2017), then contrast-transfer function (CTF) parameters estimation was done using CTFFIND4 (Rohou & Grigorieff, 2015). For each sample, the following number of micrographs were collected: Dhx29 WT - 32,069; Dhx29-K663A - 42,025 ; Dhx29- E730Q - 18,479; Dhx29-ΔN - 26,242.

### Cryo-EM data processing

For the Dhx29 wild-type dataset, 32,069 micrographs were imported for particle picking using the Relion (v5.0.0) Laplacian-of-Gaussian autopicker (Burt *et al*, 2024; Scheres, 2012). A total of 1,688,737 particles were picked and extracted at a box size of 700 pixels, 4 times binned, then imported in cryoSPARC (v4.6.0) for 2D classification (Punjani *et al*, 2017). After curation and selection of good 2D class averages, 1,322,112 particles were used for ab-initio reconstruction and homogeneous refinement. The aligned particles were re-imported into Relion and used for 3D classification. This resulted in classes containing mainly Dhx29-bound particles (POST 80S with P/P tRNA and PRE/hybrid with A/P-tRNA and P/E-tRNAs) and a smaller fraction of Dhx29-unbound particles (60S, empty 80S and some PRE/hybrid 80S). The Dhx29-bound POST 80S particles showed additional clustering depending on the presence of an A-site mRNA hairpin. For further processing, two classes showing the strongest densities of Dhx29 on POST and PRE/hybrid 80S were further processed by focused 3D classifications in Relion (Appendix Fig. S1A). For the Dhx29-POST 80S complexes, a final number of 56,757 particles were selected after focused 3D classification on the dsRBD region, extracted at a box size of 700 pixels, non-binned, then used for homogeneous refinement in cryoSPARC, resulting in a map with overall resolution of 2.4 Å (Appendix Fig. S1B, D, F, H). For the Dhx29-PRE/hybrid 80S complexes, a focused classification step based on the Dhx29 region separated two conformations of the helicase based on the 40S rotation (Appendix Fig. S1A). A final number of 201,634 particles from the most abundant class were then extracted at a box size of 700 pixels, non-binned, and used for homogenous refinement in cryoSPARC, leading to a map with overall resolution of 2.2 Å (Appendix Fig. S1C, E, G, I).

For the combined data processing, each of the four datasets (Dhx29 WT, K663A, E730Q and ΔN) was independently filtered to obtain datasets of more homogenous data quality. Micrographs were selected based on an estimated resolution cutoff of 4 Å, a defocus range of 0.5 to 2.0 μm, maximum relative astigmatism of 1.5%, and a figure of merit (FOM) cutoff of 0. Following filtering, the curated micrographs from each dataset were then randomly divided into subsets of 10,000 micrographs. One subset of 10,000 micrographs was selected from each of the four samples and used as the starting dataset for the combined processing, resulting in a total of 40,000 micrographs. This dataset was imported in Relion and the Laplacian-of-Gaussian autopicker was used to pick 2,281,556 particles. The particles were extracted at a box size of 600 pixels, 6 times binned, then imported into cryoSPARC for 2D classification. After manual curation of good 2D class averages, a total of 1,636,914 particles were used for ab-initio and homogenous refinement. The aligned particles were re-extracted at a box size of 600 pixels, 4 times binning, then 3D classified in Relion into 7 different classes of translating ribosomes, some containing no Dhx29 density (13% of particles), others containing weakly or strongly bound Dhx29 to both POST 80S (containing P/P-tRNA) and PRE/hybrid 80S (containing A/P and P/E-tRNAs) (Appendix Fig. S2A), respectively.

For the Dhx29-bound POST 80S complexes, 3D classes containing a strong Dhx29 density were selected for further processing in Relion. After one focused 3D sorting on the region comprising Dhx29 helicase and UBA-RWD modules, 202,245 particles were selected as having a strong Dhx29 density. Out of these particles, after another focused 3D sorting on the dsRBD, a final number of 57,631 particles showed strong Dhx29 dsRBD bound to an mRNA hairpin in the ribosomal A-site. These particles were extracted at a box size of 640 pixels, non-binned and used for homogenous refinement, as well al local refinements of the Dhx29 alone, resulting in an overall resolution of 2.5 Å for the full reconstruction and a resolution of 3.6 Å for the locally refined Dhx29 region (Appendix Fig. S2B,C,H,J; Appendix Table S1). The locally refined maps were fitted back into the homogenous refined map and a composite map was generated in ChimeraX (Meng *et al*, 2023). Furthermore, these particles were used for a focused classification on the mRNA hairpin in cryoSPARC, resulting in different densities showing dsRBD engaging long, consensus or mixed length mRNA HPs (Appendix Fig. S2A). Additionally, out of the 202,245 particles showing a strong Dhx29-bound POST 80S, 56,766 particles were sorted by focused cryoSPARC 3D classification as having a strong density for a neighboring ribosome. After particle extraction at a larger box size (1,000 pixels, 5 times decimated) and the removal of particles positioned at the edges of the micrographs, a total of 40,370 particles were used in cryoSPARC for a homogenous refinement of the entire collided ribosome map. Additionally, the same particles at a box size of 640 pixels, non-binned, centered on the stalled ribosome were used for a homogenous refinement resulting in a 2.6 Å map (Appendix Fig. S2A).

For the Dhx29-bound PRE/hybrid 80S complexes, 3D classes containing densities for Dhx29 (either weak or strong) were selected for further processing. After a series of focused 3D classification steps in Relion and cryoSPARC, 4 classes emerged, having different occupancy levels for the Dxh29 density or different rotations of the 40S. A total of 329,132 particles with the strongest Dhx29 density were combined, re-extracted at a box size of 640 pixels, non-binned, and used for homogenous refinement in cryoSPARC. The resulting map had an overall resolution of 2.1 Å (Appendix Fig. S2I,K). Several local refinements were also performed in cryoSPARC to improve the local resolution of the Dhx29 density, 40S head and body (Appendix Fig. S2D-G). Notably, the Dhx29 local refinement was improved and reached a resolution of 3.1 Å (Appendix Fig. S2G, see also Appendix Table S1). The locally refined maps were fitted back into the homogenous refined map and a composite map was generated in ChimeraX (v1.9) (Meng *et al*., 2023).

### Model building

For the POST-state Dhx29-bound 80S, the leading 80S of PDB:9F9S (Kim *et al*, 2024) and for Dhx29 the respective predicted model from the AF2 database (Fleming *et al*., 2025; Jumper *et al*., 2021) (AF- Q06698-F1) were used as starting models. The N- and C-terminal parts of the helicase core as well as the N-terminal helix together with the dsRBD were rigid-body docked individually and then manually adjusted. The mRNA and P-site tRNA from PDB:9F9S were manually adjusted. They serve as placeholders since the cryo-EM map represents a mixture of different mRNAs and codon positions. In the same manner, the A-site hairpin in the model only serves as a representative placeholder since it is composed of a mixture of different stem-loops (see Appendix Fig. S2 and Fig. EV4B). For the hairpin, an AF3 (Abramson *et al*, 2024) model derived from a previously reported stable stem-loop (Doma & Parker, 2006; Hosoda *et al*, 2003) that served as a template to build a 22 nt long mRNA stretch that forms a double-stranded stem that is capped by a GAAAC loop forming a GNRA tetraloop motif. For the PRE-state Dhx29-bound 80S, the collided 80S of PDB:9F9S was used as a starting model. The N- and C-terminal core parts from the Dhx29 AF2 database prediction (Fleming *et al*., 2025; Jumper *et al*., 2021) (AF-Q06698-F1) were rigid-body fitted individually and manually adjusted. Amino acid side chains for the helicase were removed where local resolution did not allow modelling. The mRNA and tRNA from PDB:9F9S were manually adjusted and serve as placeholders since the cryo-EM map represents a mixture that does not allow unambiguous identification.

All manual adjustments were done in Coot (v0.9.8.96) (Emsley *et al*, 2010) and Phenix (v2.1-6048) (Liebschner *et al*, 2019) was used for real-space refinements of both POST- and hybrid/PRE-state models.

### Selective ribosome profiling (Ribo-seq)

*DHX29*-FTpA and *dhx29*Δ strains were grown in YPD medium at 30°C and 125 rpm to a final OD_600_ of about 2.2. Cells were harvested by centrifugation and grinded using a SPEX 6970EFM Freezer/Mill. Cell powder was stored at -80°C until further use. The Dhx29-FTpA sample was purified as described above. Selective ribo-seq has been performed as previously described (Müller *et al*, 2023; Wagner *et al*, 2022). RNA concentrations of cleared lysates were measured using an Implen NP80 and 1 μL of RNase I (Ambion) was added per 120 μg of RNA and lysates were digested for 5 min on ice. For input control and DHX29 deletion ribo-seq, the digested lysate was loaded onto 800 μL of a 1 M sucrose cushion in gradient buffer (20 mM HEPES/KOH pH 7.4, 150 mM KOAc, 5 mM MgCl₂, 1 mM DTT, protease inhibitor mix). Cushions were centrifuged in a TLA-120.2 rotor (Beckman Coulter) at 434,900 x g for 1 h at 4 °C. Supernatant fractions were removed and ribosome pellets were directly resuspended in TRIzol (Thermo Fisher Scientific). RNA was extracted using the Direct-zol RNA Miniprep Kit (Zymo). Footprint size selection and subsequent library preparation were performed as previously described (Wagner *et al*., 2022).

### Visualization

Cryo-EM maps and molecular models were visualized using ChimeraX (v1.9) (Meng *et al*., 2023).

## Data availability

Cryo-EM maps and molecular models generated in this study were deposited at the Electron Microscopy Database (EMDB) and the Protein Data Bank (PDB). They are accessible via the following codes: EMD-58765 (https://www.ebi.ac.uk/emdb/EMD-58765) and 32AZ (https://doi.org/10.2210/pdb32az/pdb) (Dhx29 bound yeast 80S ribosome in POST state); EMD-58766 (https://www.ebi.ac.uk/emdb/EMD-58766) and 32BA (https://doi.org/10.2210/pdb32ba/pdb) (Dhx29 bound yeast 80S ribosome in PRE/hybrid state); EMD-58759 (https://www.ebi.ac.uk/emdb/EMD-58759) (Dhx29 bound to POST state 80S ribosome, 80S consensus); EMD-58760 (https://www.ebi.ac.uk/emdb/EMD-58760) (Dhx29 bound to POST state 80S ribosome, Dhx29 local refinement); EMD-58761 (https://www.ebi.ac.uk/emdb/EMD-58761) (Dhx29 bound to PRE/hybrid state 80S ribosome, 80S consensus); EMD-58762 (https://www.ebi.ac.uk/emdb/EMD-58762) (Dhx29 bound to PRE/hybrid state 80S ribosome, 40S body local refinement); EMD-58763 (https://www.ebi.ac.uk/emdb/EMD-58763) (Dhx29 bound to PRE/hybrid state 80S ribosome, 40S head local refinement); EMD-58764 (https://www.ebi.ac.uk/emdb/EMD-58764) (Dhx29 bound to PRE/hybrid state 80S ribosome, Dhx29 local refinement).

All sequencing data has been uploaded to the GEO database and is accessible under the identifier: GSE343842.

## Author contributions

Conceptualization: RB, MT

Data Curation: LC, TD, MBDM, OB, TB, MT

Formal Analysis: LC, TD, MBDM, TB, MT

Funding Acquisition: MBDM, RB

Investigation: LC, TD, MBDM, MT

Methodology: LC, TD, MBDM, MT

Project Administration: RB, MT

Resources: RB

Software: N/A

Supervision: MT, RB

Validation: LC, TD, MBDM, TB, MT, RB

Visualization: LC, TD, MBDM, TB

Writing (original draft): LC, TD, TB, MBDM, MT, RB

Writing (review & editing) : N/A

## Acknowledgements

We thank Susanne Rieder and Charlotte Ungewickell for technical assistance with cryo-EM data collection. This study was supported by grants from the ERC (ADG 885711 Human-Ribogenesis) and DFG (BE1814/20-1, BE1814/22-1) to RB and (MU 5391/1-1) to MBDM; LC was supported by the International Max Planck School for Molecules of Life (IMPRS-ML); TD was supported by the Graduate School of Quantitative Biosciences Munich (QBM); RB was funded by DFG - Project-ID 533767322 - EXC 3113/1, Cluster for Nucleic Acid Sciences and Technologies - NUCLEATE.

